# Associations between brain structure and sleep patterns across adolescent development

**DOI:** 10.1101/2021.01.05.424689

**Authors:** Maria Jalbrzikowski, Rebecca Hayes, Kathleen E. Scully, Peter L. Franzen, Brant P. Hasler, Greg J. Siegle, Daniel J. Buysse, Ron E. Dahl, Erika E. Forbes, Cecile D. Ladouceur, Dana L. McMakin, Neal D. Ryan, Jennifer S. Silk, Tina R. Goldstein, Adriane M. Soehner

## Abstract

**Importance:** Structural brain maturation and sleep are complex processes that exhibit significant changes over adolescence and are linked to healthy physical and mental development. The precise timing and magnitude of these changes influence function throughout the lifespan. However, the relationships between gray matter structure and sleep patterns during adolescence are not fully understood. A detailed characterization of brain-sleep associations during this sensitive period is crucial for understanding factors contributing to optimal neurodevelopmental trajectories in adolescence.

**Objective:** To investigate whether sleep-gray matter relationships are developmentally-invariant (i.e., stable across age) or developmentally-specific (i.e., only present during discrete time windows) from late childhood through young adulthood.

**Setting:** The Neuroimaging and Pediatric Sleep Databank was constructed from 8 research studies conducted at the University of Pittsburgh between 2009 and 2020.

**Participants:** The final sample consisted of 240 participants without current psychiatric diagnoses (9-25 years), and with good quality sleep tracking and structural MRI (sMRI) data.

**Design:** Participants completed a sMRI scan and 5-7 days of wrist actigraphy to assess naturalistic sleep. We examined cross-sectional associations between sMRI measures and sleep patterns, as well as the effects of age, sex, and their interaction with sMRI measures on sleep.

**Main Outcome(s) and Measure(s):** Using Freesurfer software, we extracted cortical thickness and subcortical volumes from T1-weighted MRI. Sleep patterns (duration, timing, continuity, regularity) were estimated from wrist actigraphy.

**Results:** Shorter sleep duration, later sleep timing, and poorer sleep continuity were associated with a stable pattern of thinner cortex and altered subcortical volumes in diverse brain regions across adolescence. In a discrete subset of regions (e.g., posterior cingulate), thinner cortex was associated with these sleep patterns from late childhood through early-to-mid adolescence, but not in late adolescence and young adulthood.

**Conclusions and Relevance:** In childhood and adolescence, developmentally-invariant and developmentally-specific associations exist between sleep patterns and gray matter structure, in a wide array of brain regions linked to many sensory, cognitive, and emotional processes. Sleep intervention during specific developmental periods could potentially promote healthier neurodevelopmental outcomes.

**KEY POINTS:** *Question:* Does age modulate associations between gray matter structure and actigraphic sleep patterns across adolescent development?

*Findings:* This cross-sectional study reports stable associations between regional gray matter structure and shorter duration, later timing, and poorer continuity of sleep from ages 9 to 25 years-old, as well as developmentally-specific associations that are present only from late childhood to early-to-mid adolescence.

*Meaning:* Stronger coupling of gray matter and sleep patterns from late childhood to early-to-mid adolescence potentially implicates this discrete developmental window as a period of vulnerability to adverse sleep-brain interactions. Sleep intervention during this developmental stage may support healthier neurodevelopmental trajectories.

## INTRODUCTION

Structural brain maturation and sleep are complex processes that exhibit significant changes during adolescent development. The precise timing and amount of these changes in youths likely influences multiple adult outcomes. Optimal sleep and brain maturation are each known to influence adolescent health and functioning, including academic/vocational achievement, mental health, and/or risk behaviors^1–10^. However, relationships between gray matter structure and sleep patterns over adolescence are not fully understood; furthermore, it is unknown whether these relationships vary as a function of age. A detailed characterization of brain-sleep relationships in adolescence is important for understanding factors contributing to optimal neurodevelopmental trajectories during this sensitive period.

Many brain regions implicated in cognitive and emotional outcomes show a protracted developmental course through adolescence^11–15^, indicating that periods of heightened plasticity also come with greater vulnerability^15,16^. Cortical thickness usually peaks by age 9-10 and then decreases until early adulthood, particularly in frontal, parietal, and temporal regions^4,12,17–23^. Most subcortical regions increase in volume until ∼14-15 years, with growth plateauing afterwards^13,24,25^. Deviations from these normative trajectories may increase vulnerability to diverse negative outcomes, including poorer academic performance, mental health difficulties, and/or risky behaviors.

During adolescence, brain structural maturation is accompanied by multiple cognitive, behavioral, and emotional changes, including changes in sleep. Adolescence is characterized by a circadian phase delay and reduced homeostatic sleep drive, contributing to later sleep timing^26,27^. These biological shifts converge with psychosocial and behavioral factors (e.g., school start times, peer socializing) to result in insufficient sleep and, at times, poorer sleep regularity or continuity^26,27^. Disruptions to the timing, duration, continuity, and regularity of sleep predict and track with the severity of adverse cognitive and emotional outcomes (e.g., poor school performance, depression, substance use)^28–32^.

Developmental shifts in sleep characteristics may possess reciprocal relationships with brain structural maturation^33–36^, ultimately influencing diverse outcomes. While sleep serves multiple purposes, one such function is to support synaptic plasticity and reorganization of brain circuitry^24^. Sleep disruption was originally considered a *consequence* of brain structural abnormalities; however, recent animal data indicate that sleep disruption during periods of heightened developmental plasticity also *cause* deviations in brain maturation^37–39^. These translational studies imply stronger brain-sleep relationships in certain developmental windows^37,40^. Yet, in humans it is unknown whether brain-sleep relationships are stable across adolescent development (i.e., developmentally-invariant relationships) or only occur during a discrete window of development (i.e., developmentally-specific relationships). Developmentally-specific brain-sleep relationships could inform the optimal timing of brain and/or sleep-based interventions that promote healthier neurodevelopmental outcomes. Several initial reports have identified ties between diverse gray matter structures and sleep in pediatric populations^41–48^.

However, developmentally-specific relationships have not been examined and these studies have been restricted to retrospective self-report or lab-based sleep measures that do not reflect usual sleep. An important next step is to evaluate how brain structure relates to objective, ecologically-valid sleep patterns (as captured by wrist actigraphy) through a developmental lens.

To address these open questions, we created the Neuroimaging and Pediatric Sleep (NAPS) Databank, a large, harmonized cross-sectional databank comprised of healthy children, adolescents, and young adults (ages 9-25yr). We estimated sleep from wrist actigraphy and sMRI measures from T1-weighted MRI. Given that a wide array of sMRI measures have been associated with sleep, we conducted data-driven regularized regression analyses, to test many potential predictors while minimizing the issues of predictor inter-correlation and multiple comparisons. We explored developmentally-invariant and developmentally-specific associations between sMRI measures (subcortical volume, cortical thickness) and core sleep dimensions (sleep duration, timing, continuity, regularity). Because there are important sex differences in sleep and brain development^17,49–54^, we also explored the interaction between self-reported sex and neuroimaging measures on sleep outcomes.

## METHODS

### Participants

The initial NAPS databank includes a total of 307 participants drawn from eight University of Pittsburgh studies conducted between the years of 2009 to 2020. Studies were considered for inclusion in NAPS if they included: a) actigraphic sleep monitoring; b) a sMRI scan; c) a baseline assessment reflecting naturalistic sleep; and d) participants aged 8-30 years-old (inclusive). Participant-level inclusion criteria were: a) 8-25 years-old; b) absence of current psychiatric diagnosis based on clinical interview (i.e., KSADS, SCID); c) no current psychotropic or hypnotic medication use; d) ≥5 days of good quality actigraphic sleep monitoring composed of both weekday and weekend days; e) good quality MRI scan. Demographics of the final analytic sample of N=240 are described in **Table 1**. Demographics by protocol are reported in **eTable 1** and reasons for participant exclusion are documented in the **eMethods**.

**Table 1:** NAPS sample characteristics

| Variable | Mean or n (sd or %) |
| --- | --- |
| Sample N | 240 |
| Age (Years) | 18.13 (5.26) |
| Self-reported Sex |  |
| Female | 133 (55%) |
| Male | 107 (45%) |
| Ethnicity |  |
| Non-Hispanic | 223 (93%) |
| Hispanic | 15 (6%) |
| Missing | 2 (1%) |
| Race |  |
| White | 170 (71%) |
| Black | 40 (17%) |
| Asian | 11 (5%) |
| Multiple | 16 (7%) |
| Unknown/Missing | 3 (1%) |
| Wrist Actigraph Type |  |
| AMI Octagonal MotionLogger | 36 (15%) |
| PR/MiniMitter Actiwatch64 | 25 (10%) |
| PR Actiwatch2 | 113 (47%) |
| PR Spectrum Series | 66 (28%) |
| Tracking Days | 6.59 (0.84) |
| Weekdays | 4.51 (0.92) |
| Weekend Days | 2.08 (0.52) |
| Season |  |
| Spring | 43 (18%) |
| Summer | 100 (42%) |
| Fall | 52 (22%) |
| Winter | 45 (19%) |
| Sleep Duration (min) | 417.96 (63.52) |
| Wake After Sleep Onset (minutes) | 56.19 (26.83) |
| Midsleep (minutes from midnight) | 289.99 (76.59) |
| Midsleep Variability (minutes) | 63.66 (44.06) |

### Neuroimaging Methods and Outcomes

Please see **eTable 2** for sMRI protocol parameters. We used the FreeSurfer analysis software^55–58^ (v6.0) to extract measures of cortical thickness (Desikan-Killiany atlas^59^, n=34 measures) and subcortical volume (aseg.mgz atlas, n=8 measures) averaged across two hemispheres. We implemented a quality assessment pipeline developed by and used for the Enhancing Neuroimaging Genetics through Meta-Analysis consortium^60–70^. An automated MRIQC T1w-classifier determined individual scan quality based on a reference template^71^. We adjusted neuroimaging data for scanner protocol effects with ComBat^72,73^.

### Wrist Actigraphy

Actigraphy is a well-validated and widely-used tool for objectively assessing naturalistic sleep in children, adolescents, and adults^74–76^. Participants continuously wore wrist actigraphs on their non-dominant wrist during a monitoring period of 5 or more consecutive days^77^. **eTable 1** describes the number of participants who wore watches from Philips Respironics (PR; Actiwach-64, Actiwatch2, Spectrum series) or Ambulatory Monitoring, Inc. (AMI; Basic Octagonal Motionlogger). Wrist activity was sampled in 1-minute intervals (epochs). Participants were asked to indicate via button press the start and end of each sleep interval.

We estimated sleep from wrist actigraphy using a combination of validated brand-specific sleep algorithms (PR Medium Threshold; AMI Sadeh) and standardized visual editing procedures^78–80^. Trained scorers blinded to neuroimaging data manually identified rest intervals based on a combination of event markers indicated by participants and clear changes in activity and (if available) environmental light level recorded by the device. Brand-specific sleep scoring algorithms estimated sleep within each rest interval^74,75,79,81–83^. We implemented additional semi-automated quality assurance procedures using in-house R scripts, including identification of the main rest interval (defined as the longest rest interval each day), removal of invalid sleep intervals containing ≥1 hour of off-wrist time or recording errors^79,84^, time adjustment for daylight savings time, and final visual inspection of sleep intervals on raster plots.

### Sleep Outcomes

Primary actigraphy sleep outcomes were based on the main rest interval. We selected four sleep outcomes corresponding to key dimensions of sleep health^85^: sleep duration (total sleep time in minutes), timing (midpoint between sleep onset and offset in minutes from midnight), continuity (minutes awake after sleep onset; WASO), and regularity (intra-individual standard deviation of midpoint in minutes). The first three outcomes were averaged over the 5-7 tracking days most proximal to their MRI scan; regularity was calculated from the available days of recording. Sleep variables were natural log transformed to normalize distributions.

### Statistical Analyses

We first conducted general additive models to confirm that the four sleep outcomes showed age-associated patterns consistent with prior research (**eFigure 1**). We observed the characteristic decline in sleep duration, delay in sleep timing, and increased sleep variability over adolescent development. Sleep continuity did not vary with age.

We were interested in developmentally-invariant effects (i.e., main effects) of neuroimaging measures on the four sleep outcomes, as well as developmentally-specific effects (i.e., interactions between age and neuroimaging measures). Due to the large number of and multicollinearity amongst neuroimaging measures, we used regularized regression^86^ to identify non-zero predictors associated with sleep outcomes. We used the R package, Group-Lasso-INTERaction-NET (glinternet^87,88^) to examine main effects of structural neuroimaging measures, as well as their interaction with age and sex, for each sleep variable. Only potential interactions between non-zero main effects are considered. We included multiple actigraphy covariates (i.e., tracking days, season, ratio of weekday to weekend days, actigraph model) as potential predictors in the models. **eTable 3** contains the full list of 48 predictors. We repeated 10-fold cross validation 100 times, using the penalty parameter (λ) one standard deviation away from the minimal cross-validation error. The final model was the model was selected most often during this procedure. Regularized regression selects variables based on minimizing error in the model as opposed to statistical significance as in standard regression. Thus, p-values are not reported for non-zero coefficients.

Non-zero predictors selected by group-lasso models were entered into linear regression models, as in prior reports^89,90^. R-squared was computed to estimate variance explained by the full model as well as groups of predictors (i.e., demographics, neuroimaging measures, actigraphy covariates). We assessed non-zero interactions between age and neuroimaging predictors with the Johnson-Neyman technique, which obtains parameter estimates and points of significance from the interaction between two continuous variables^91–93^. Non-zero interactions between sex and neuroimaging predictors were probed by comparing estimated marginal means^94^.

## RESULTS

All neuroimaging measures, and their interactions with age and sex, selected as non-zero predictors of sleep outcomes are reported in **Table 2**. Non-zero actigraphy covariates (e.g., season, actigraph type) are reported in **eTable 4**.

**Table 2.** Main effects and interactions between age, sex, and neuroimaging measures on actigraphic sleep dimensions. Model weights are reported as standardized regression coefficients.

| **A. Sleep Duration (Total Sleep Time)** |  |  |
| --- | --- | --- |
| *Type of Effect* | *Variable* | *Model Weight* |
| Demographic Variable Main Effects | Sex | 0.0403 |
|  | Age | -0.0732 |
| Subcortical Volume Main Effects | Pallidum | -0.0122 |
|  | Hippocampus | -0.0529 |
|  | Amygdala | -0.0032 |
|  | Lateral Ventricles | 0.0221 |
| Cortical Thickness Main Effects | Medial Orbitofrontal Cortex | 0.1090 |
|  | Parahippocampal Cortex | 0.0022 |
|  | Posterior Cingulate | 0.0576 |
|  | Isthmus Cingulate | 0.0196 |
|  | Superior Parietal Cortex | 0.0067 |
|  | Cuneus | 0.0277 |
| Sex Interactions | Sex x Parahippocampal Cortex | 0.0054 |
|  | Sex by Posterior Cingulate Cortex | -0.0219 |
| Age Interactions | Age x Lateral Ventricles | 0.0234 |
|  | Age x Cuneus | -0.0332 |
|  | Age x Superior Parietal Cortex | -0.0072 |
| Variance Accounted for by demographic measures only: $R^2=0.22$ | | |
| Variance Accounted for by neuroimaging and demographic measures, and their interactions: $R^2=0.25$ | | |
| **B. Sleep Timing (Midsleep)** |  |  |
| *Type of Effect* | *Variable* | *Model Weight* |
| Demographic Variable Main Effects | Sex | -0.0601 |
|  | Age | 0.1315 |
| Subcortical Volume Main Effects | Thalamus | -0.0009 |
|  | Pallidum | 0.0120 |
|  | Lateral Ventricles | 0.0057 |
| Cortical Thickness Main Effects | Medial Orbitofrontal Cortex | -0.0002 |
|  | Pars Orbitalis | -0.0136 |
|  | Rostral Middle Frontal Cortex | -0.0200 |
|  | Posterior Cingulate Cortex | -0.0089 |
|  | Superior Parietal Cortex | -0.0051 |
|  | Lateral Occipital Cortex | -0.1115 |
| Sex Interactions | Sex x Lateral Ventricles | 0.0342 |
|  | Sex x Thalamus | 0.0010 |
| Age Interactions | Age x Pallidum | -0.0431 |
|  | Age x Medial Orbitofrontal Cortex | 0.0002 |
|  | Age x Pars Orbitalis | 0.0187 |
|  | Age x Rostral Middle Frontal Cortex | 0.0267 |
|  | Age x Posterior Cingulate Cortex | 0.0189 |
| Variance Accounted for by demographic measures only: $R^2=0.10$ | | |
| Variance Account for by neuroimaging and demographic measures, and their interactions: $R^2=0.20$ | | |

| **C. Sleep Continuity (WASO)** |  |  |
| --- | --- | --- |
| *Type of Effect* | *Variable* | *Model Weight* |
| Demographic Variable Main Effects | Sex | -0.0444 |
|  | Age | 0.0244 |
| Subcortical Volume Main Effects | Thalamus | 0.0065 |
|  | Pallidum | 0.0260 |
|  | Caudate | 0.0083 |
| Cortical Thickness Main Effects | Entorhinal Cortex | 0.0009 |
|  | Parahippocampal Cortex | -0.0192 |
|  | Middle Temporal Cortex | -0.0086 |
|  | Precentral Cortex | -0.0283 |
|  | Superior Parietal Cortex | -0.0243 |
|  | Lateral Occipital Cortex | -0.0149 |
| Sex Interactions | Sex x Caudate | -0.0117 |
|  | Sex x Entorhinal Cortex | -0.0021 |
|  | Sex x Precentral Cortex | -0.0151 |
| Age Interactions | Age x Parahippocampal Cortex | 0.0533 |
|  | Age x Superior Parietal Cortex | 0.0622 |
| Variance Accounted for by demographic measures only: $R^2=0.05$ | | |
| Variance Account for by neuroimaging and demographic measures, and their interactions: $R^2=0.16$ | | |

### Sleep Duration (Total Sleep Time)

The main effects of neuroimaging measures, age, sex, and their respective interactions accounted for 25% of the total variance in sleep duration (**Table 2A**). Shorter sleep duration was associated with older age and males had shorter sleep duration in comparison to females.

We observed several developmentally-invariant relationships between brain structure and sleep duration. From 9-25 years old, greater volume in the pallidum, hippocampus, and amygdala was associated with shorter sleep duration. Additionally, thinner medial orbitofrontal and isthmus (posterior) cingulate cortices were associated shorter sleep duration. Thinner cortex in the posterior cingulate was associated with shorter sleep duration in both sexes, but there was a stronger relationship in males. Conversely, thinner parahippocampal cortex and shorter sleep duration were associated in females, but not males.

We also found developmentally-specific relationships between gray matter structure and sleep duration (**Figure 2A**). In late childhood through middle adolescence, thinner cortex in the cuneus (9-17.3 years) and superior parietal regions (9-16.0 years) was associated with shorter sleep duration; however, this relationship was not observed at older ages. From 21.9-25.9 years old, greater lateral ventricle volume was associated with longer sleep duration.

**Figure 1.**
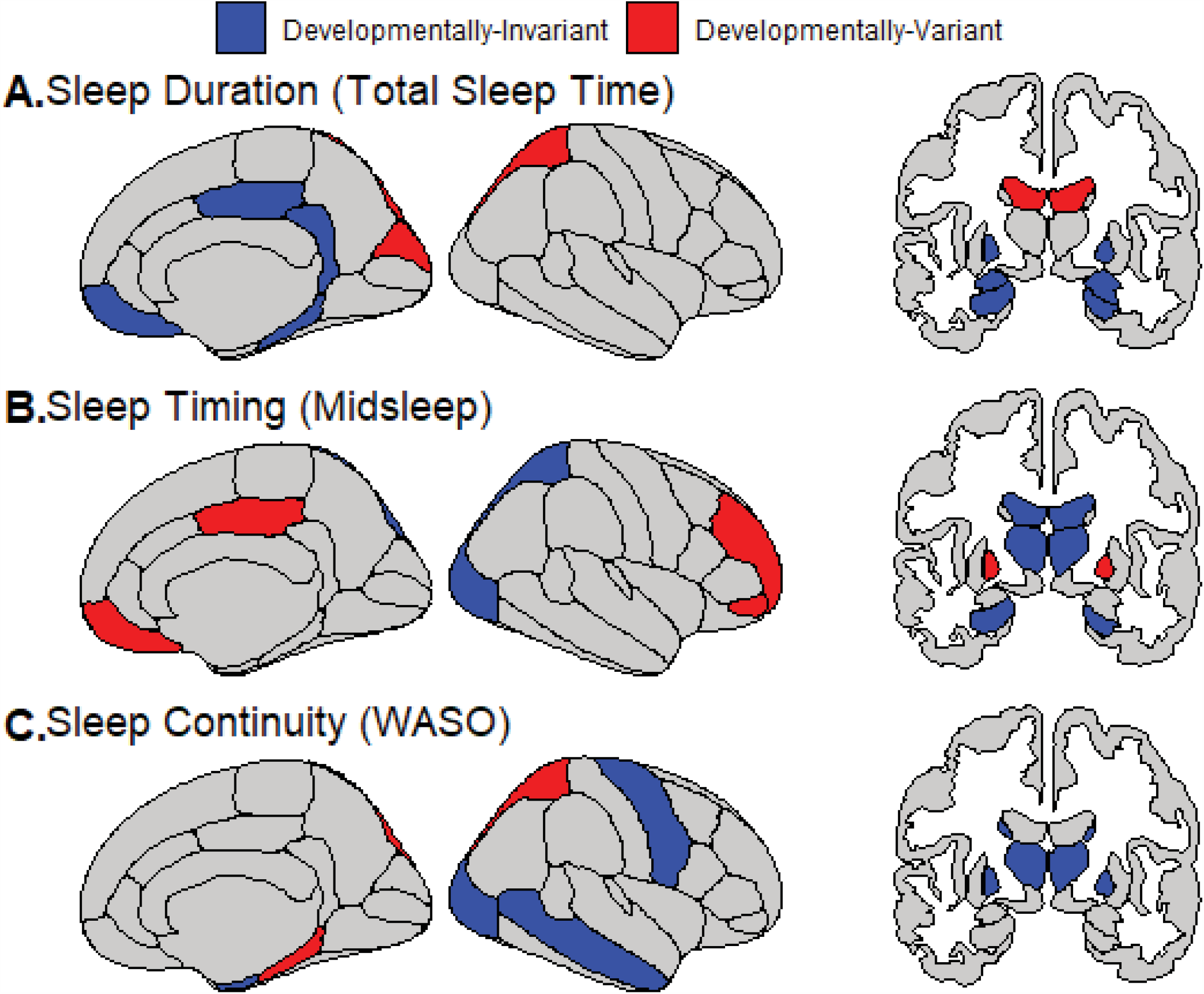
Relationships between sleep and gray matter (cortical thickness, subcortical volume) that are developmentally-invariant (i.e., stable across age) or developmentally-specific (i.e., only present during discrete time windows) from late childhood through young adulthood.

**Figure 2.**
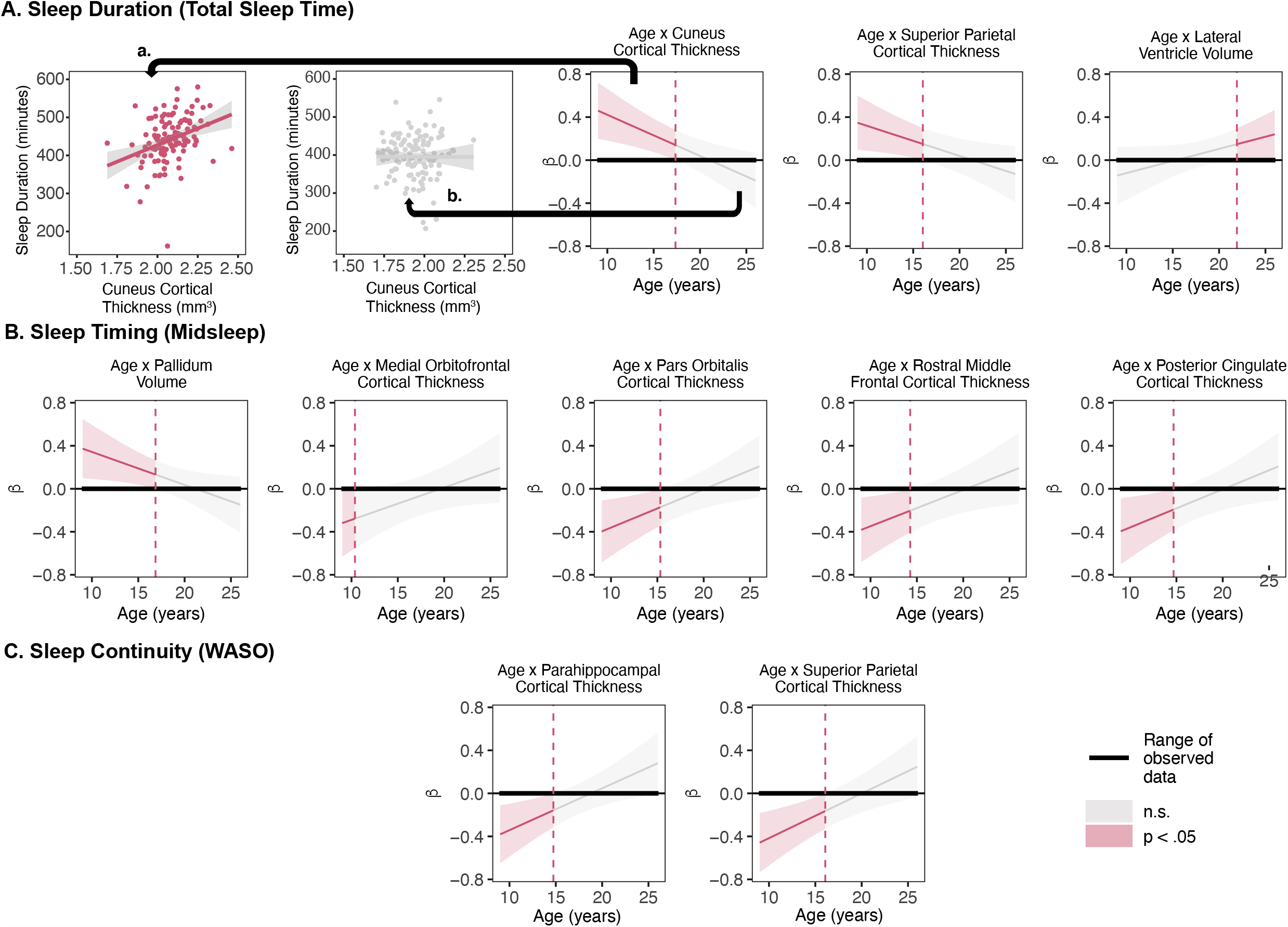
Johnsyon-Neyman plots of age by neuroimaging measure interactions on sleep dimensions (A. duration, B. timing, and C. continuity). A statistically significant relationship between age and the neuroimaging measures (p<.05) is represented by the red color. Non-significant relationships are represented by the gray color. To aid in the interpretation of the plots, we provide one example of the age by cuneus cortical thickness interaction on sleep duration. a. From 9-17.3 years old, thicker cuneus cortex is associated with longer sleep duration (*r*=0.33, *p*=1.0×10^−4^). b. From 17.4-25.9 years old, this relationship is not present (r=-0.003, p=0.97).

### Sleep Timing (Midsleep)

The main effects of neuroimaging measures, age, and their interactions accounted for 20% of the variance in midsleep (**Table 2B**). Midsleep was later in males and among older participants.

Developmentally-invariant relationships were identified for several brain regions. Specifically, lower thalamus volume was associated with later midsleep; this was relationship driven by males. In females only, greater lateral ventricle volume was associated with later midsleep. Thinner superior parietal and lateral occipital cortices were associated with later sleep timing.

Developmentally-specific relationships were also observed between neuroimaging measures and sleep timing (**Figure 2B**). From late childhood through middle adolescence, thinner cortex in the pars orbitalis (9-15.2 years), rostral middle frontal (9-14.1 years), and posterior cingulate regions (9-14.5 years) was associated with later midsleep. Thinner medial orbitofrontal cortex in late childhood (9-10 years) was also associated with later midsleep. Greater pallidum volume was associated with later midsleep only from ages 9 to 16.8 years. **Sleep Continuity (WASO)**

The combined effects of neuroimaging measures, age, sex, and their interactions accounted for 16% of the variance in sleep continuity (**Table 2C**). WASO was longer among older participants and in females.

With regard to developmentally-invariant relationships, greater palladium and thalamus volume was associated with greater WASO. Thinner cortex in middle temporal, precentral, and lateral occipital regions was associated with greater WASO. Greater precentral and entorhinal cortical thickness was associated with greater WASO in females.

Thinner parahippocampal (9-14.6 years) and superior parietal cortices (9-16.0 years) were associated with greater WASO from late childhood to mid-adolescence, but not in older adolescents and young adults (**Figure 2C**).

### Sleep Regularity (Midsleep Variability)

Regularized regression did not identify any nonzero predictors of midsleep regularity.

## DISCUSSION

Using a large sample of typical adolescent development (9.0-25.9 years), we identified developmentally-invariant and developmentally-specific relationships between gray matter structure and naturalistic sleep patterns. Shorter sleep duration, later sleep timing, and poorer sleep continuity — all of which are associated with adverse health outcomes — were associated with a stable pattern of thinner cortex and altered subcortical volumes in diverse brain regions over adolescent development. In discrete regions, developmentally-specific relationships were also observed. In these regions, thinner cortex from late childhood through early-to-mid adolescence — a pattern associated with accelerated maturation — was associated with less optimal sleep, but these relationships were not detected in late adolescence and young adulthood. Our results provide a novel view of brain-sleep structure relationships within brain structures implicated in a wide array of cognitive, emotional, and psychological processes over adolescent development^2,95–100^.

### Cortical thickness in a diverse set of brain regions show developmentally-invariant relationships with sleep

Across adolescent development, thinner cortex in frontal, temporal, parietal, and visual processing areas was associated with shorter sleep duration, later sleep timing, and longer time awake after sleep onset. These brain regions are implicated in salience detection (pars orbitalis), motor function (precentral), memory (entorhinal, middle temporal), and attention and visuospatial perception (superior parietal cortex, lateral occipital)^101^. Given that sleep is associated with diverse range of mental, cognitive and physical health outcomes in adolescence^1–10^, it is reasonable that naturalistic sleep is related to brain structure in regions that support multiple functions. Some of these relationships were modulated by self-reported sex, consistent with reported sex differences in sleep patterns and brain development^17,49–54^. Future studies should also examine the extent to sex effects may be better explained by pubertal maturation.

### Increased cortical thickness was associated with healthier sleep patterns from late childhood to middle adolescence

This is the first study, to our knowledge, to demonstrate that brain structure is related to individual differences in naturalistic sleep patterns at different ages, from late childhood through adulthood. Thicker cortex in multiple brain regions was associated with “healthier” sleep (as indicated by longer, more continuous, and earlier sleep) during late childhood and early adolescence. These findings, in conjunction with other work^102^, present the possibility that biological factors exert differential influences on behavior at distinct points in development. Accelerated cortical thinning/growth patterns in discrete brain regions could contribute to disruptions in sleep characteristics during late childhood and early adolescence, but not during other periods. Alternatively, disruptions in the typical age-related changes in sleep could lead to accelerating cortical thinning, particularly during this late childhood-early adolescence age range, but not during others. Multiple neurobiological mechanisms likely underlie individual differences in cortical thickness. Cortical thinning is traditionally believed to be caused by synaptic pruning, a re-wiring of synapses^103,104^. Translational models find that, in mice, synaptic pruning is *higher* during sleep than wakefulness in adolescents, but not adults^105^. More recent data suggest that age-associated changes in cortical thickness may also be driven by white matter maturational processes, i.e. myelination^106^. Sleep disruption is detrimental to the formation and maintenance of myelin in murine models^107,108^. Future longitudinal within-person investigations, particularly during late childhood and early adolescence, will be necessary to disentangle the directionality and neurobiological mechanisms of relationships between sleep, cortical thickness measures, and white matter integrity.

### Unexpected relationships between poorer sleep and larger subcortical volumes

Surprisingly, in many cases, we also discovered that *larger* subcortical (i.e., hippocampal, amygdala, thalamus, and caudate) volumes are associated with more disrupted sleep patterns. One possibility is that exposure to sleep disruption at certain developmental stages may be correlated with or cause *accelerated* subcortical growth patterns, akin to the acceleration-deceleration hypothesis of chronic stress and neurodevelopment^109–111^. Importantly, this result stands in contrast with prior research showing lower subcortical gray matter volumes in relation to poor sleep^46^ and mental health conditions^60,112,113^. Thus, replication of these findings, as well as work examining the relationship between structural brain measures and sleep, needs to be further explored in informative subgroups such as individuals with mental disorders.

We also observed subcortical volume-sleep relationships in the expected direction. In females, larger lateral ventricle volume was associated with shorter sleep duration and later midsleep. Greater ventricle size has been linked to serious mental health conditions, including schizophrenia^114^. Furthermore, study of older adults also found longitudinal reduction in sleep duration corresponded to ventricular expansion over the follow-up period^115^.

### Implications for optimal timing and targets for sleep intervention

If sleep patterns prove to be a causal contributor to individual differences in sMRI measures, our findings have the potential to inform developmentally-sensitive optimization of evidence-based behavioral sleep interventions^116^. As an example, both shorter sleep duration and later sleep timing were associated with thinner cortex in default mode network (DMN) regions (medial orbitofrontal and posterior cingulate cortices), a neural signature tied to outcomes such as depression, insomnia, and poor cognitive function^98,117^. DMN cortical thickness and sleep duration relationships were developmentally-invariant. However, DMN cortical thickness-sleep timing association were only present in late childhood/mid-adolescence. Thus, a sleep treatment geared toward promoting healthy DMN-relevant outcomes should include sleep extension regardless of age but also advance sleep timing in late childhood and early/mid adolescence. Taken as a whole, our findings suggest that sleep interventions, particularly in late childhood through mid-adolescence, may be advantageous for neurodevelopment and thus downstream effects on psychological well-being.

## Limitations

Our sample, while representative of the Pittsburgh Metropolitan area, was limited in its racial and ethnic diversity, factors which contribute to individual differences in brain structure and sleep^28,118^. Although we adjusted for salient actigraphy covariates, actigraphy brand differences may have contributed noise in our data that was not captured by covarying for watch type in our models. Because our analyses were cross-sectional across a range of ages, rather than longitudinal within participants, it is unclear whether sleep patterns are a cause, correlate, or consequence of gray matter structure. Future, prospective longitudinal studies are necessary to disambiguate causal relationships between sleep and sMRI measures, and assess relationships between within-subject trajectories of sleep and brain development.

## Conclusions & Future Directions

We found compelling and novel evidence for developmentally-invariant and developmentally-specific associations between sMRI measures and sleep across adolescent development. We plan to build on these findings and examine how individual differences in neuroimaging and sleep measures may identify youth at high-risk for developing adverse cognitive, mental, and physical outcomes.

## Author Contributions

Drs. Soehner and Jalbrzikowski had full access to all the data and take responsibility for the integrity of the data and the accuracy of the data analysis.

*Concept and design of NAPS databank:* Jalbrzikowski, Soehner

*Concept, design, and funding for original research studies:* Franzen, Hasler, Siegle, Buysse, Dahl, Forbes, Ladouceur, McMakin, Ryan, Silk, Goldstein, Soehner

*Acquisition, processing, or interpretation of data:* All authors.

*Statistical analysis:* Jalbrzikowski, Soehner

*Drafting of the manuscript:* Jalbrzikowski, Soehner

*Critical revision of the manuscript for important intellectual content:* All authors.

## Previous Presentation

These results have not previously been presented or submitted for publication.

## REFERENCES

1. Pehlivanova M, Wolf DH, Sotiras A, et al. Diminished Cortical Thickness Is Associated with Impulsive Choice in Adolescence. J Neurosci. 2018;38(10):2471–2481. doi:10.1523/JNEUROSCI.2200-17.2018

2. Burgaleta M, Johnson W, Waber DP, Colom R, Karama S. Cognitive ability changes and dynamics of cortical thickness development in healthy children and adolescents. Neuroimage. 2014;84:810–819. doi:10.1016/j.neuroimage.2013.09.038

3. Foland-Ross LC, Sacchet MD, Prasad G, Gilbert B, Thompson PM, Gotlib IH. Cortical thickness predicts the first onset of major depression in adolescence. Int J Dev Neurosci. 2015;46:125–131. doi:10.1016/j.ijdevneu.2015.07.007

4. Tamnes CK, Ostby Y, Fjell AM, Westlye LT, Due-Tønnessen P, Walhovd KB. Brain maturation in adolescence and young adulthood: regional age-related changes in cortical thickness and white matter volume and microstructure. Cereb Cortex. 2010;20(3):534–548. doi:10.1093/cercor/bhp118

5. Bos MGN, Peters S, van de Kamp FC, Crone EA, Tamnes CK. Emerging depression in adolescence coincides with accelerated frontal cortical thinning. J Child Psychol Psychiatry. 2018;59(9):994–1002. doi:10.1111/jcpp.12895

6. Oostermeijer S, Whittle S, Suo C, et al. Trajectories of adolescent conduct problems in relation to cortical thickness development: a longitudinal MRI study. Transl Psychiatry. 2016;6(6):e841. doi:10.1038/tp.2016.111

7. Meruelo AD, Jacobus J, Idy E, Nguyen-Louie T, Brown G, Tapert SF. Early adolescent brain markers of late adolescent academic functioning. Brain Imaging Behav. 2019;13(4):945–952. doi:10.1007/s11682-018-9912-2

8. Dewald JF, Meijer AM, Oort FJ, Kerkhof GA, Bögels SM. The influence of sleep quality, sleep duration and sleepiness on school performance in children and adolescents: A metaanalytic review. Sleep Med Rev. 2010;14(3):179–189. doi:10.1016/j.smrv.2009.10.004

9. McKnight-Eily LR, Eaton DK, Lowry R, Croft JB, Presley-Cantrell L, Perry GS. Relationships between hours of sleep and health-risk behaviors in US adolescent students. Prev Med. 2011;53(4-5):271–273. doi:10.1016/j.ypmed.2011.06.020

10. Tarokh L, Saletin JM, Carskadon MA. Sleep in adolescence: physiology, cognition and mental health. Neurosci Biobehav Rev. 2016;70:182–188. doi:10.1016/j.neubiorev.2016.08.008

11. Giedd JN, White SL, Celano M. Structural Magnetic Resonance Imaging of Typical Pediatric Brain Development. In: Charney D, Nestler E, eds. Neurobiology of Mental Illness. Oxford University Press; 2011:1209–1217. doi:10.1093/med/9780199798261.003.0074

12. Giedd JN, Raznahan A, Alexander-Bloch A, Schmitt E, Gogtay N, Rapoport JL. Child Psychiatry Branch of the National Institute of Mental Health Longitudinal Structural Magnetic Resonance Imaging Study of Human Brain Development. Neuropsychopharmacology. 2015;40(1):43–49. doi:10.1038/npp.2014.236

13. Raznahan A, Shaw PW, Lerch JP, et al. Longitudinal four-dimensional mapping of subcortical anatomy in human development. Proc Natl Acad Sci U S A. 2014;111(4):1592–1597. doi:10.1073/pnas.1316911111

14. Giedd JN, Blumenthal J, Jeffries NO, et al. Brain development during childhood and adolescence: a longitudinal MRI study. Nature Neuroscience. 1999;2(10):861–863. doi:10.1038/13158

15. Paus T, Keshavan M, Giedd JN. Why do many psychiatric disorders emerge during adolescence? Nat Rev Neurosci. 2008;9(12):947–957.

16. Dahl RE. Adolescent brain development: a period of vulnerabilities and opportunities. Keynote address. Ann N Y Acad Sci. 2004;1021:1–22. doi:10.1196/annals.1308.001

17. Vijayakumar N, Allen NB, Youssef G, et al. Brain development during adolescence: A mixed-longitudinal investigation of cortical thickness, surface area, and volume. Hum Brain Mapp. 2016;37(6):2027–2038. doi:10.1002/hbm.23154

18. Tamnes CK, Herting MM, Goddings A-L, et al. Development of the Cerebral Cortex across Adolescence: A Multisample Study of Inter-Related Longitudinal Changes in Cortical Volume, Surface Area, and Thickness. J Neurosci. 2017;37(12):3402–3412. doi:10.1523/JNEUROSCI.3302-16.2017

19. Mills KL, Goddings A-L, Herting MM, et al. Structural brain development between childhood and adulthood: Convergence across four longitudinal samples. Neuroimage. 2016;141:273–281.

20. Sowell ER, Peterson BS, Thompson PM, Welcome SE, Henkenius AL, Toga AW. Mapping cortical change across the human life span. Nat Neurosci. 2003;6(3):309–315. doi:10.1038/nn1008

21. Sowell ER, Thompson PM, Leonard CM, Welcome SE, Kan E, Toga AW. Longitudinal mapping of cortical thickness and brain growth in normal children. J Neurosci. 2004;24(38):8223–8231. doi:10.1523/JNEUROSCI.1798-04.2004

22. Gogtay N, Giedd JN, Lusk L, et al. Dynamic mapping of human cortical development during childhood through early adulthood. Proc Natl Acad Sci U S A. 2004;101(21):8174– 8179.

23. Shaw P, Greenstein D, Lerch J, et al. Intellectual ability and cortical development in children and adolescents. Nature. 2006;440(7084):676–679. doi:10.1038/nature04513

24. Tononi G, Cirelli C. Sleep and the price of plasticity: from synaptic and cellular homeostasis to memory consolidation and integration. Neuron. 2014;81(1):12–34. doi:10.1016/j.neuron.2013.12.025

25. Puentes-Mestril C, Aton SJ. Linking Network Activity to Synaptic Plasticity during Sleep: Hypotheses and Recent Data. Front Neural Circuits. 2017;11. doi:10.3389/fncir.2017.00061

26. Crowley SJ, Wolfson AR, Tarokh L, Carskadon MA. An update on adolescent sleep: New evidence informing the perfect storm model. J Adolesc. 2018;67:55–65. doi:10.1016/j.adolescence.2018.06.001

27. Crowley SJ, Van Reen E, LeBourgeois MK, et al. A longitudinal assessment of sleep timing, circadian phase, and phase angle of entrainment across human adolescence. PLoS One. 2014;9(11):e112199. doi:10.1371/journal.pone.0112199

28. Grandner MA. Sleep, Health, and Society. Sleep Med Clin. 2017;12(1):1–22. doi:10.1016/j.jsmc.2016.10.012

29. Spira AP, Chen-Edinboro LP, Wu MN, Yaffe K. Impact of sleep on the risk of cognitive decline and dementia. Curr Opin Psychiatry. 2014;27(6):478–483. doi:10.1097/YCO.0000000000000106

30. Matricciani L, Bin YS, Lallukka T, et al. Rethinking the sleep-health link. Sleep Health. 2018;4(4):339–348. doi:10.1016/j.sleh.2018.05.004

31. Dolsen MR, Asarnow LD, Harvey AG. Insomnia as a transdiagnostic process in psychiatric disorders. Curr Psychiatry Rep. 2014;16(9):471. doi:10.1007/s11920-014-0471-y

32. Soehner AM, Kaplan KA, Harvey AG. Insomnia comorbid to severe psychiatric illness. Sleep Med Clin. 2013;8(3):361–371. doi:10.1016/j.jsmc.2013.04.007

33. Kayser MS, Biron D. Sleep and Development in Genetically Tractable Model Organisms. Genetics. 2016;203(1):21–33. doi:10.1534/genetics.116.189589

34. Muehlroth BE, Werkle-Bergner M. Understanding the interplay of sleep and aging: Methodological challenges. Psychophysiology. 2020;57(3). doi:10.1111/psyp.13523

35. Fontanellaz-Castiglione CE, Markovic A, Tarokh L. Sleep and the adolescent brain. Current Opinion in Physiology. 2020;15:167–171. doi:10.1016/j.cophys.2020.01.008

36. Cirelli C, Tononi G. Cortical development, electroencephalogram rhythms, and the sleep/wake cycle. Biol Psychiatry. 2015;77(12):1071–1078. doi:10.1016/j.biopsych.2014.12.017

37. Kayser MS, Yue Z, Sehgal A. A Critical Period of Sleep for Development of Courtship Circuitry and Behavior in Drosophila. Science. 2014;344(6181):269–274. doi:10.1126/science.1250553

38. Seugnet L, Suzuki Y, Donlea JM, Gottschalk L, Shaw PJ. Sleep Deprivation During Early-Adult Development Results in Long-Lasting Learning Deficits in Adult Drosophila. Sleep. 2011;34(2):137–146.

39. Frank MG, Issa NP, Stryker MP. Sleep enhances plasticity in the developing visual cortex. Neuron. 2001;30(1):275–287. doi:10.1016/s0896-6273(01)00279-3

40. Novati A, Hulshof HJ, Koolhaas JM, Lucassen PJ, Meerlo P. Chronic sleep restriction causes a decrease in hippocampal volume in adolescent rats, which is not explained by changes in glucocorticoid levels or neurogenesis. Neuroscience. 2011;190:145–155. doi:10.1016/j.neuroscience.2011.06.027

41. Goldstone A, Willoughby AR, de Zambotti M, et al. The mediating role of cortical thickness and gray matter volume on sleep slow-wave activity during adolescence. Brain Struct Funct. 2018;223(2):669–685. doi:10.1007/s00429-017-1509-9

42. Cheng W, Rolls E, Gong W, et al. Sleep duration, brain structure, and psychiatric and cognitive problems in children. Mol Psychiatry. Published online February 3, 2020. doi:10.1038/s41380-020-0663-2

43. Urrila AS, Artiges E, Massicotte J, et al. Sleep habits, academic performance, and the adolescent brain structure. Sci Rep. 2017;7. doi:10.1038/srep41678

44. Kurth S, Ringli M, Geiger A, LeBourgeois M, Jenni OG, Huber R. Mapping of Cortical Activity in the First Two Decades of Life: A High-Density Sleep Electroencephalogram Study. J Neurosci. 2010;30(40):13211–13219. doi:10.1523/JNEUROSCI.2532-10.2010

45. Sung D, Park B, Kim S-Y, et al. Structural Alterations in Large-scale Brain Networks and Their Relationship with Sleep Disturbances in the Adolescent Population. Sci Rep. 2020;10(1):3853. doi:10.1038/s41598-020-60692-1

46. Taki Y, Hashizume H, Thyreau B, et al. Sleep duration during weekdays affects hippocampal gray matter volume in healthy children. Neuroimage. 2012;60(1):471–475.

47. Lapidaire W, Urrila AS, Artiges E, et al. Sleep, regional grey matter volumes, and psychological functioning in adolescents. bioRxiv. Published online November 30, 2019. doi:10.1101/645184

48. Buchmann A, Ringli M, Kurth S, et al. EEG sleep slow-wave activity as a mirror of cortical maturation. Cereb Cortex. 2011;21(3):607–615. doi:10.1093/cercor/bhq129

49. Sowell ER, Trauner DA, Gamst A, Jernigan TL. Development of cortical and subcortical brain structures in childhood and adolescence: a structural MRI study. Dev Med Child Neurol. 2002;44(1):4–16. doi:10.1017/s0012162201001591

50. Koolschijn PCMP, Crone EA. Sex differences and structural brain maturation from childhood to early adulthood. Dev Cogn Neurosci. 2013;5:106–118. doi:10.1016/j.dcn.2013.02.003

51. Zhang B, Wing Y-K. Sex differences in insomnia: a meta-analysis. Sleep. 2006;29(1):85–93. doi:10.1093/sleep/29.1.85

52. Santhi N, Lazar AS, McCabe PJ, Lo JC, Groeger JA, Dijk D-J. Sex differences in the circadian regulation of sleep and waking cognition in humans. Proc Natl Acad Sci U S A. 2016;113(19):E2730–2739. doi:10.1073/pnas.1521637113

53. Giedd JN, Castellanos FX, Rajapakse JC, Vaituzis AC, Rapoport JL. Sexual dimorphism of the developing human brain. Prog Neuropsychopharmacol Biol Psychiatry. 1997;21(8):1185–1201. doi:10.1016/s0278-5846(97)00158-9

54. Lenroot RK, Gogtay N, Greenstein DK, et al. Sexual dimorphism of brain developmental trajectories during childhood and adolescence. Neuroimage. 2007;36(4):1065–1073. doi:10.1016/j.neuroimage.2007.03.053

55. Dale AM, Fischl B, Sereno MI. Cortical surface-based analysis. I. Segmentation and surface reconstruction. Neuroimage. 1999;9(2):179–194.

56. Fischl B, Sereno MI, Dale AM. Cortical surface-based analysis. II: Inflation, flattening, and a surface-based coordinate system. Neuroimage. 1999;9(2):195–207.

57. Fischl B, Dale AM. Measuring the thickness of the human cerebral cortex from magnetic resonance images. Proc Natl Acad Sci U S A. 2000;97(20):11050–11055. doi:10.1073/pnas.200033797

58. Fischl B, Salat DH, Busa E, et al. Whole brain segmentation: automated labeling of neuroanatomical structures in the human brain. Neuron. 2002;33(3):341–355.

59. Desikan RS, Segonne F, Fischl B, et al. An automated labeling system for subdividing the human cerebral cortex on MRI scans into gyral based regions of interest. Neuroimage. 2006;31(3):968–980.

60. Schmaal L, Veltman DJ, TGM van Erp, et al. Subcortical brain alterations in major depressive disorder: findings from the ENIGMA Major Depressive Disorder working group. Mol Psychiatry. 2016;21(6):806–812. doi:10.1038/mp.2015.69

61. Hibar DP, Westlye LT, Doan NT, et al. Cortical abnormalities in bipolar disorder: an MRI analysis of 6503 individuals from the ENIGMA Bipolar Disorder Working Group. Mol Psychiatry. 2018;23(4):932–942.

62. TGM van Erp, Walton E, Hibar DP, et al. Cortical Brain Abnormalities in 4474 Individuals With Schizophrenia and 5098 Control Subjects via the Enhancing Neuro Imaging Genetics Through Meta Analysis (ENIGMA) Consortium. Biological Psychiatry. 2018;84(9):644–654. doi:10.1016/j.biopsych.2018.04.023

63. van Rooij D, Anagnostou E, Arango C, et al. Cortical and Subcortical Brain Morphometry Differences Between Patients With Autism Spectrum Disorder and Healthy Individuals Across the Lifespan: Results From the ENIGMA ASD Working Group. Am J Psychiatry. 2018;175(4):359–369. doi:10.1176/appi.ajp.2017.17010100

64. Hoogman M, Bralten J, Hibar DP, et al. Subcortical brain volume differences in participants with attention deficit hyperactivity disorder in children and adults: a cross-sectional mega-analysis. Lancet Psychiatry. 2017;4(4):310–319.

65. Sun D, Ching CRK, Lin A, et al. Large-scale mapping of cortical alterations in 22q11.2 deletion syndrome: Convergence with idiopathic psychosis and effects of deletion size. Mol Psychiatry. Published online June 13, 2018. doi:10.1038/s41380-018-0078-5

66. Kong X-Z, Mathias SR, Guadalupe T, et al. Mapping cortical brain asymmetry in 17,141 healthy individuals worldwide via the ENIGMA Consortium. Proc Natl Acad Sci U S A. 2018;115(22):E5154–E5163. doi:10.1073/pnas.1718418115

67. Boedhoe PSW, Schmaal L, Abe Y, et al. Cortical Abnormalities Associated With Pediatric and Adult Obsessive-Compulsive Disorder: Findings From the ENIGMA Obsessive-Compulsive Disorder Working Group. Am J Psychiatry. 2018;175(5):453–462. doi:10.1176/appi.ajp.2017.17050485

68. Whelan CD, Altmann A, Botía JA, et al. Structural brain abnormalities in the common epilepsies assessed in a worldwide ENIGMA study. Brain. 2018;141(2):391–408.

69. Nunes A, Schnack HG, Ching CRK, et al. Using structural MRI to identify bipolar disorders - 13 site machine learning study in 3020 individuals from the ENIGMA Bipolar Disorders Working Group. Mol Psychiatry. Published online August 2018.

70. Thompson PM, Stein JL, Medland SE, et al. The ENIGMA Consortium: large-scale collaborative analyses of neuroimaging and genetic data. Brain Imaging Behav. 2014;8(2):153–182. doi:10.1007/s11682-013-9269-5

71. Esteban O, Birman D, Schaer M, Koyejo OO, Poldrack RA, Gorgolewski KJ. MRIQC: Advancing the automatic prediction of image quality in MRI from unseen sites. PLoS One. 2017;12(9):e0184661. doi:10.1371/journal.pone.0184661

72. Fortin J-P, Parker D, Tunç B, et al. Harmonization of multi-site diffusion tensor imaging data. Neuroimage. 2017;161:149–170.

73. Fortin J-P, Cullen N, Sheline YI, et al. Harmonization of cortical thickness measurements across scanners and sites. Neuroimage. 2018;167:104–120.

74. Quante M, Kaplan ER, Cailler M, et al. Actigraphy-based sleep estimation in adolescents and adults: a comparison with polysomnography using two scoring algorithms. Nat Sci Sleep. 2018;10:13–20. doi:10.2147/NSS.S151085

75. Meltzer LJ, Walsh CM, Traylor J, Westin AML. Direct Comparison of Two New Actigraphs and Polysomnography in Children and Adolescents. Sleep. 2012;35(1):159–166. doi:10.5665/sleep.1608

76. Meltzer LJ, Wong P, Biggs SN, et al. Validation of Actigraphy in Middle Childhood. Sleep. 2016;39(6):1219–1224. doi:10.5665/sleep.5836

77. Acebo C, Sadeh A, Seifer R, et al. Estimating Sleep Patterns with Activity Monitoring in Children and Adolescents: How Many Nights Are Necessary for Reliable Measures? Sleep. 1999;22(1):95–103. doi:10.1093/sleep/22.1.95

78. Meltzer LJ, Montgomery-Downs HE, Insana SP, Walsh CM. Use of Actigraphy for Assessment in Pediatric Sleep Research. Sleep Med Rev. 2012;16(5):463–475. doi:10.1016/j.smrv.2011.10.002

79. Patel SR, Weng J, Rueschman M, et al. Reproducibility of a Standardized Actigraphy Scoring Algorithm for Sleep in a US Hispanic/Latino Population. Sleep. 2015;38(9):1497–1503. doi:10.5665/sleep.4998

80. Blackwell T, Ancoli-Israel S, Gehrman PR, Schneider JL, Pedula KL, Stone KL. Actigraphy scoring reliability in the study of osteoporotic fractures. Sleep. 2005;28(12):1599–1605. doi:10.1093/sleep/28.12.1599

81. Kanady JC, Drummond SPA, Mednick SC. Actigraphic assessment of a polysomnographic-recorded nap: a validation study. J Sleep Res. 2011;20(1 Pt 2):214–222. doi:10.1111/j.1365-2869.2010.00858.x

82. van Hees VT, Sabia S, Jones SE, et al. Estimating sleep parameters using an accelerometer without sleep diary. Sci Rep. 2018;8(1):12975. doi:10.1038/s41598-018-31266-z

83. Slater JA, Botsis T, Walsh J, King S, Straker LM, Eastwood PR. Assessing sleep using hip and wrist actigraphy. Sleep Biol Rhythms. 2015;13(2):172–180. doi:10.1111/sbr.12103

84. Feliciano EC, Rifas-Shiman S, Quante M, Redline S, Oken E, Taveras E. Chronotype, Social Jet Lag, and Cardiometabolic Risk Factors in Early Adolescence. Jama Pediatrics. 2019;173(11):1049–1057. doi:10.1001/jamapediatrics.2019.3089

85. Buysse DJ. Sleep health: can we define it? Does it matter? Sleep. 2014;37(1):9–17.

86. Tibshirani R. Regression Shrinkage and Selection Via the Lasso. Journal of the Royal Statistical Society, Series B. 1994;58:267–288.

87. Lim M, Hastie T. Learning interactions via hierarchical group-lasso regularization. J Comput Graph Stat. 2015;24(3):627–654. doi:10.1080/10618600.2014.938812

88. Lim M, Hastie T. Glinternet: Learning Interactions via Hierarchical Group-Lasso Regularization.; 2020. Accessed December 8, 2020. https://CRAN.R-project.org/package=glinternet

89. Soehner AM, Bertocci MA, Levenson JC, et al. Longitudinal associations between sleep patterns and psychiatric symptom severity in high-risk and community comparison youth. J Am Acad Child Adolesc Psychiatry. 2019;58(6):608–617. doi:10.1016/j.jaac.2018.09.448

90. Bertocci MA, Hanford L, Manelis A, et al. Clinical, cortical thickness and neural activity predictors of future affective lability in youth at risk for bipolar disorder: initial discovery and independent sample replication. Mol Psychiatry. 2019;24(12):1856–1867. doi:10.1038/s41380-018-0273-4

91. Johnson PO, Fay LC. The Johnson-Neyman technique, its theory and application. Psychometrika. 1950;15(4):349–367. doi:10.1007/BF02288864

92. Curran PJ, Bauer DJ. Testing and probing interactions in hierarchical linear growth models. Methodological issues in …. Published online 2006.

93. Preacher KJ, Curran PJ, Bauer DJ. Computational Tools for Probing Interactions in Multiple Linear Regression, Multilevel Modeling, and Latent Curve Analysis. Journal of Educational and Behavioral Statistics. 2006;31(3):437–448.

94. Lenth R, Buerkner P, Herve M, Love J, Riebl H, Singmann H. Emmeans: Estimated Marginal Means, Aka Least-Squares Means.; 2020. Accessed December 8, 2020. https://CRAN.R-project.org/package=emmeans

95. Patel P, Miles AE, Nikolova YS. Cortical thickness correlates of probabilistic reward learning in young adults. Biol Psychol. 2020;157:107975. doi:10.1016/j.biopsycho.2020.107975

96. Tamnes CK, Overbye K, Ferschmann L, et al. Social perspective taking is associated with self-reported prosocial behavior and regional cortical thickness across adolescence. Dev Psychol. 2018;54(9):1745–1757. doi:10.1037/dev0000541

97. Karama S, Colom R, Johnson W, et al. Cortical thickness correlates of specific cognitive performance accounted for by the general factor of intelligence in healthy children aged 6 to 18. Neuroimage. 2011;55(4):1443–1453. doi:10.1016/j.neuroimage.2011.01.016

98. Li Q, Zhao Y, Chen Z, et al. Meta-analysis of cortical thickness abnormalities in medication-free patients with major depressive disorder. Neuropsychopharmacology. 2020;45(4):703–712. doi:10.1038/s41386-019-0563-9

99. Brühl AB, Delsignore A, Komossa K, Weidt S. Neuroimaging in social anxiety disorder—a meta-analytic review resulting in a new neurofunctional model. Neurosci Biobehav Rev. 2014;47:260–280. doi:10.1016/j.neubiorev.2014.08.003

100. Bora E, Fornito A, Yücel M, Pantelis C. Voxelwise meta-analysis of gray matter abnormalities in bipolar disorder. Biol Psychiatry. 2010;67(11):1097–1105. doi:10.1016/j.biopsych.2010.01.020

101. Smitha K, Akhil Raja K, Arun K, et al. Resting state fMRI: A review on methods in resting state connectivity analysis and resting state networks. Neuroradiol J. 2017;30(4):305–317. doi:10.1177/1971400917697342

102. Jalbrzikowski M, Larsen B, Hallquist MN, Foran W, Calabro F, Luna B. Development of White Matter Microstructure and Intrinsic Functional Connectivity Between the Amygdala and Ventromedial Prefrontal Cortex: Associations With Anxiety and Depression. Biol Psychiatry. 2017;82(7):511–521. doi:10.1016/j.biopsych.2017.01.008

103. Huttenlocher PR. Synaptic density in human frontal cortex - developmental changes and effects of aging. Brain Res. 1979;163(2):195–205. doi:10.1016/0006-8993(79)90349-4

104. Rakic P, Bourgeois JP, Eckenhoff MF, Zecevic N, Goldman-Rakic PS. Concurrent overproduction of synapses in diverse regions of the primate cerebral cortex. Science. 1986;232(4747):232–235. doi:10.1126/science.3952506

105. Maret S, Faraguna U, Nelson AB, Cirelli C, Tononi G. Sleep and waking modulate spine turnover in the adolescent mouse cortex. Nat Neurosci. 2011;14(11):1418–1420. doi:10.1038/nn.2934

106. Natu VS, Gomez J, Barnett M, et al. Apparent thinning of human visual cortex during childhood is associated with myelination. Proc Natl Acad Sci U S A. 2019;116(41):20750–20759. doi:10.1073/pnas.1904931116

107. Bellesi M, Haswell JD, de Vivo L, et al. Myelin modifications after chronic sleep loss in adolescent mice. Sleep. 2018;41(5). doi:10.1093/sleep/zsy034

108. Ganji SK, An Z, Banerjee A, Madan A, Hulsey KM, Choi C. Measurement of regional variation of GABA in the human brain by optimized point-resolved spectroscopy at 7 T in vivo. NMR Biomed. 2014;27(10):1167–1175.

109. Tottenham N, Sheridan MA. A Review of Adversity, The Amygdala and the Hippocampus: A Consideration of Developmental Timing. Front Hum Neurosci. 2010;3. doi:10.3389/neuro.09.068.2009

110. Merz EC, Tottenham N, Noble KG. Socioeconomic Status, Amygdala Volume, and Internalizing Symptoms in Children and Adolescents. J Clin Child Adolesc Psychol. 2018;47(2):312–323. doi:10.1080/15374416.2017.1326122

111. Yoo S-S, Gujar N, Hu P, Jolesz FA, Walker MP. The human emotional brain without sleep--a prefrontal amygdala disconnect. Curr Biol. 2007;17(20):R877–878. doi:10.1016/j.cub.2007.08.007

112. TGM van Erp, Hibar DP, Rasmussen JM, et al. Subcortical brain volume abnormalities in 2028 individuals with schizophrenia and 2540 healthy controls via the ENIGMA consortium. Molecular Psychiatry. 2016;21(4):547–553. doi:10.1038/mp.2015.63

113. Logue MW, SJH van Rooij, Dennis EL, et al. Smaller Hippocampal Volume in Posttraumatic Stress Disorder: A Multisite ENIGMA-PGC Study: Subcortical Volumetry Results From Posttraumatic Stress Disorder Consortia. Biol Psychiatry. 2018;83(3):244–253. doi:10.1016/j.biopsych.2017.09.006

114. Chung Y, Haut KM, He G, et al. Ventricular enlargement and progressive reduction of cortical gray matter are linked in prodromal youth who develop psychosis. Schizophr Res. 2017;189:169–174. doi:10.1016/j.schres.2017.02.014

115. Lo JC, Loh KK, Zheng H, Sim SKY, Chee MWL. Sleep Duration and Age-Related Changes in Brain Structure and Cognitive Performance. Sleep. 2014;37(7):1171–1178. doi:10.5665/sleep.3832

116. Blake MJ, Sheeber LB, Youssef GJ, Raniti MB, Allen NB. Systematic Review and Metaanalysis of Adolescent Cognitive-Behavioral Sleep Interventions. Clin Child Fam Psychol Rev. 2017;20(3):227–249. doi:10.1007/s10567-017-0234-5

117. Suh S, Kim H, Dang-Vu TT, Joo E, Shin C. Cortical Thinning and Altered Cortico-Cortical Structural Covariance of the Default Mode Network in Patients with Persistent Insomnia Symptoms. Sleep. 2016;39(1):161–171. doi:10.5665/sleep.5340

118. LeWinn KZ, Sheridan MA, Keyes KM, Hamilton A, McLaughlin KA. Sample composition alters associations between age and brain structure. Nat Commun. 2017;8. doi:10.1038/s41467-017-00908-7

