## supplemental text & tables for "Associations between brain structure and sleep patterns across adolescent development"

### eMethods

**Participants.** Of the total 307 cases in NAPS, cases were excluded based on: enrollment in multiple protocols (n=3), age > 25 years-old (n=18); presence of a psychiatric diagnosis (n=21); poor quality or insufficient sleep tracking (n=7); or poor quality MRI (n=18).

### eResults

**eTable 1. Demographic and actigraph characteristics by study**

| Study | N | Female (%) | Age Range | Age Mean (SD) | Enrollment Year (Min-Max) | Actigraph (Brand-Model) |
|---|---|---|---|---|---|---|
| ACRES | 22 | 11 (50.0%) | 13.3-17.8 | 15.5 (1.27) | 2013-2017 | Philips-Spectrum |
| BASS | 21 | 16 (76.2%) | 13.65-23.02 | 19.43 (2.76) | 2012-2013 | Philips-Actiwatch2 |
| BEST | 18 | 9 (50.0%) | 14.1-18.53 | 15.76 (1.16) | 2017-2019 | Philips-Spectrum |
| CATS | 36 | 20 (55.6%) | 9.16-14.53 | 11.49 (1.56) | 2009-2013 | AMI-Motionlogger |
| PAN2 | 41 | 22 (53.7%) | 11.41-14.78 | 13.18 (0.97) | 2012-2014 | Philips-Actiwatch2 |
| SCARAB | 25 | 16 (64.0%) | 18.73-22.75 | 21.12 (1.21) | 2014-2017 | Philips-Spectrum |
| SDMR | 25 | 14 (56.0%) | 18.69-25.52 | 22.9 (1.68) | 2009-2012 | Philips/MiniMitter-Actiwatch64 |
| SIRA | 51 | 26 (51.0%) | 19.27-30.09 | 24.53 (2.94) | 2014-2016 | Philips-Actiwatch2 |

**eTable 2. MRI acquisition parameters by study**

| Study | Scanner Manufacturer | Tesla Strength | Scanner Model | Repetition Time (ms) | Echo Time (ms) | Inversion Time (ms) | Flip Angle | Voxel Size (mm) | Phase Field of View |
|---|---|---|---|---|---|---|---|---|---|
| ACRES | Siemens | 3 | Trio Tim | 2.2 | 0.00331 | 1 | 9 | 1 | 75 |
| ACRES | Siemens | 3 | Biograph mMR | 2.2 | 0.00335 | 1 | 9 | 1 | 75 |
| BASS | Siemens | 3 | Trio Tim | 2.2 | 0.00331 | 1 | 9 | 1 | 75 |
| BEST | Siemens | 3 | Prisma | 5.0 | 0.00297 | .7/2.5 | 4/5 | 1 | 93.8 |
| CATS | Siemens | 3 | Trio Tim | 2.1 | 0.00343 | 1.05 | 8 | 1 | 81.25 |
| PAN2 | Siemens | 3 | Trio Tim | 2.2 | 0.00331 | 1 | 9 | 1 | 75 |
| SCARAB | Siemens | 3 | Trio Tim | 1.5 | 0.00319 | 0.8 | 8 | 1 | 100 |
| SDMR | Siemens | 3 | Trio Tim | 2.2 | 0.00331 | 1 | 9 | 1 | 75 |
| SIRA | Siemens | 3 | Trio Tim | 2.1 | 0.00331 | 1.05 | 8 | 1 | 81.25 |

| **eTable 3. List of all predictors entered into the group-lasso model.** |
|---|
| **Demographic measures**<br>Age, Sex |
| **Neuroimaging measures**<br>*Global measures:* Estimated intracranial volume, mean cortical thickness<br>*Cortical thickness:* banks of the superior temporal sulcus, caudal anterior cingulate, caudal middle frontal, cuneus, entorhinal, frontal pole, fusiform, inferior parietal, inferior temporal, insula, isthmus cingulate, lateral occipital lateral orbitofrontal, lingual, medial orbitofrontal, middle temporal, paracentral, parahippocampal, pars opercularis, pars orbitalis, pars triangularis, pericalcarine, postcentral, posterior cingulate, precentral, precuneus, rostral anterior cingulate, rostral middle frontal, superior frontal, superior parietal, superior temporal, supramarginal, temporal pole, transverse temporal<br>*Subcortical volumes:* nucleus accumbens, amygdala, caudate, hippocampus, lateral ventricles, pallidum, putamen, thalamus |
| **Actigraphy covariates**<br>Wrist actigraph brand-model, number of tracking days, weekend:weekday ratio, season |

**eTable 4. Group-lasso actigraphy covariates selected as non-zero predictors.**

| Variable | Model Weight |
|---|---|
| **A. Sleep Duration** | |
| Wrist Actigraph (Brand-Model) | 0.031 |
| Tracking Days | -0.144 |
| Weekend:Weekday Ratio | -0.008 |
| Sex x Weekend:Weekday Ratio | 0.013 |
| Age x Wrist Actigraph (Brand-Model) | -0.037 |
| **B. Midsleep Average** | |
| Wrist Actigraph (Brand-Model) | -0.024 |
| Season | 0.009 |
| Sex x Wrist Actigraph (Brand-Model) | 0.023 |
| Sex x Season | 0.008 |
| Age x Wrist Actigraph (Brand-Model) | 0.018 |
| Age x Season | 0.053 |
| **C. Wake After Sleep Onset** | |
| Season | 0.011 |
| Wrist Actigraph (Brand-Model) | -0.001 |
| Sex x Season | 0.006 |
| Age x Wrist Actigraph (Brand-Model) | 0.052 |

**eFigure 1. Change in actigraphic sleep patterns as a function of age**

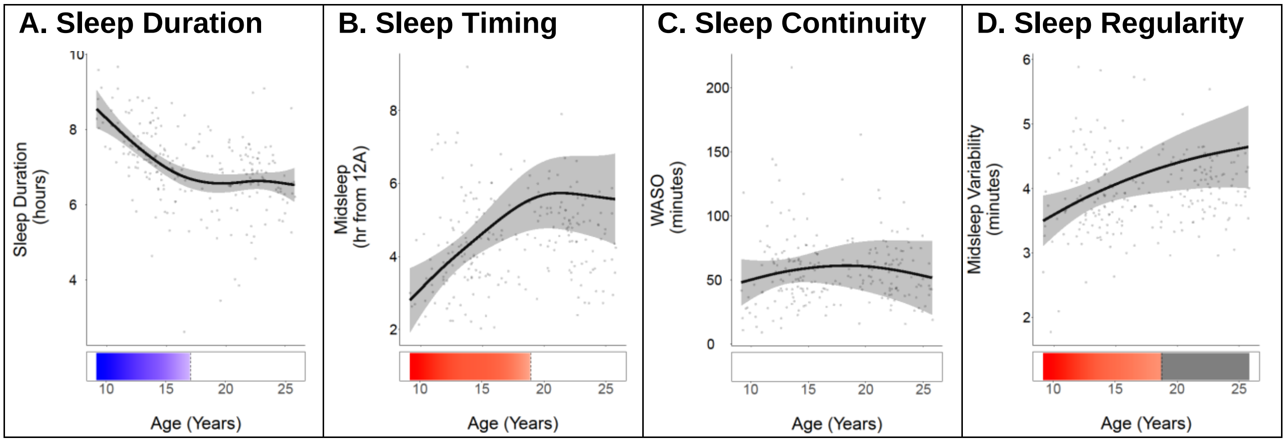


Note: Generalized additive models adjusted for sex, study, number of tracking days, weekday to weekend day ratio, season, and actigraph model. Red reflects period of significant positive slope (p<.05), blue indicates range of significant negative slope (p<.05), white/grey reflects non significance.
